# Simulations for Designing and Interpreting Intervention Trials in Infectious Diseases

**DOI:** 10.1101/198051

**Authors:** M. Elizabeth Halloran, Kari Auranen, Sarah Baird, Nicole E. Basta, Steve Bellan, Ron Brookmeyer, Ben Cooper, Victor DeGruttola, James Hughes, Justin Lessler, Eric T. Lofgren, Ira M. Longini, Jukka-Pekka Onnela, Berk Özler, George Seage, Thomas A. Smith, Alessandro Vespignani, Emilia Vynnycky, Marc Lipsitch

**Affiliations:** Vaccine and Infectious Disease Division, Fred Hutchinson Cancer Research Center, Seattle WA, USA; Department of Biostatistics, University of Washington, Seattle WA, USA; Department of Mathematics and Statistics, University of Turku, Turku, Finland; Institute for International Economic Policy, The George Washington University, Washington DC, USA; Department of Epidemiology, University of Minnesota, Minneapolis MN, USA; Department of Epidemiology and Biostatistics, University of Georgia, Athens GA, USA; Department of Biostatistics, UCLA, Los Angeles CA, USA; Mahidol Oxford Tropical Medicine Research Unit, Bangkok, Thailand; Department of Biostatistics, Harvard T.H. Chan School of Public Health, Boston MA, USA; Department of Epidemiology, Johns Hopkins School of Public Health, Baltimore MD, USA; Paul G. Allen School for Global Animal Health, Washington State University, Pullman WA, USA; Department of Biostatistics, University of Florida, Gainesville FL, USA; World Bank, Washington DC, USA; Department of Epidemiology, Harvard T.H. Chan School of Public Health, Boston MA, USA; Swiss Tropical and Public Health Institute, Basel, Switzerland; Network Science Institute, Northeastern University, Boston MA, USA; London School of Hygiene and Tropical Medicine, London, UK

**Keywords:** clinical trial design, infectious diseases, simulations, vaccine

## Abstract

Here we urge the adoption of a new paradigm for the design and interpretation of intervention trials in infectious diseases, particularly in emerging infectious disease, that more accurately reflects the dynamics of the transmission process. Interventions in infectious diseases can have indirect effects on those not receiving the intervention as well as direct effects on those receiving the intervention. Combinations of interventions can have complex interactions at the population level. These often cannot be adequately addressed with standard study designs and analytic methods. Simulations can help to accurately represent transmission dynamics in an increasingly complex world which is critical for proper trial design and interpretation. Some ethical aspects of a trial can also be quantified using simulations. After a trial has been conducted, simulations can be used to explore possible explanations for the observed effects. A great deal is to be gained through a multidisciplinary approach that builds collaborations among experts in infectious disease dynamics, epidemiology, statistical science, economics, simulation methods and the conduct of clinical trials.

---

Designing intervention trials in infectious diseases poses many challenges. First, in many infectious diseases, interventions can have indirect effects on individuals not receiving the intervention as well as on those receiving the intervention. Such indirect effects, sometimes called spillover effects, may affect estimation of the direct effects and are also of public health significance themselves. Second, because transmission is a nonlinear and stochastic process, outcomes in different arms of an intervention trial may be more variable than expected in a population where each individual’s outcome is statistically independent of those of other individuals. Third, heterogeneity from different sources, such as host susceptibility, pathogen variability, and exposure heterogeneity, can complicate study design. Fourth, the effects of a combination of interventions in a trial, such as vaccination and behavioral intervention, may be difficult to predict at the design phase. Other factors, including logistical complexities and ethical considerations can add to these challenges. After a trial has been conducted, interpreting unexpected trial results can be difficult.

Recently, investigators have used computer simulation to assist in the design, analysis and interpretation of randomized trials of infectious disease prevention measures to address these challenges. Here we describe these challenges in more detail and illustrate ways in which simulation can help to conduct better trials and to improve understanding of trial results. We conclude by advocating that for many infectious disease prevention trials, simulating the trial with the underlying transmission dynamics is an efficient way to compare different designs and to identify key aspects critical to its success, thereby improving the choice of design.

## 1 Challenges of Designing Intervention Studies for Infectious Diseases and the Role of Simulations

In the design phase of randomized trials of social or biomedical interventions, investigators consider options for how to conduct the trial and ultimately choose the trial population(s), the intervention or control conditions that will occur in each trial arm, the primary and secondary outcomes to be measured, and the way in which randomization will occur. For any set of such choices, a biostatistician working on the study can estimate the range of likely outcomes that could occur in trials of various sizes, then estimate the required sample size to achieve a specified power. This estimate often comes from closed-form equations that produce accurate sample size estimates under defined assumptions about the expected frequency of the outcome in the absence and presence of the intervention, and the amount of variability expected in the outcome within and between the arms of the trial. For many applications outside of infectious diseases, plausible assumptions about these quantities can be made directly based on previous trials, preclinical studies, or theoretical considerations. In particular, for noninfectious diseases, disease frequencies in participants randomized to the control group can reasonably be assumed to be similar to those in groups of individuals not receiving the intervention. Frequencies in those randomized to intervention can be assumed to be the same as those in the control group, reduced by a factor proportional to coverage and adherence of the intervention times the efficacy of the intervention. In infectious diseases, these assumptions are often violated due to the indirect effects of the intervention.

### 1.1 Complications for direct effects

In trials of vaccines and some other measures to prevent infectious diseases, interventions on each trial participant may affect the risk of the outcome on others, both participants and nonparticipants in the trial. For example, recipients of an efficacious vaccine are less likely to become infected but also may be less infectious if they are infected. Infectious diseases are an example of dependent happenings, where the frequency of the outcome depends on the number already affected, which can be changed by intervention [1]. Figure 1 illustrates some of the different effects that might occur in infectious disease interventions [2, 3]. Consider two clusters, or populations, of individuals. In one of the populations, a certain portion of individuals is vaccinated and the rest remain unvaccinated. In the other population, no one is vaccinated. The *direct effect* of vaccination in the population in which some individuals were vaccinated is defined by comparing the average outcomes in vaccinated individuals with the average outcomes in unvaccinated individuals. The *indirect effects* are defined as a contrast between the average outcomes in unvaccinated individuals in the population with vaccination and the average outcomes of unvaccinated individuals in the unvaccinated population. The *total effects* are defined by comparing the average outcomes in the vaccinated individuals in the vaccinated population to the average outcomes in the unvaccinated individuals in the unvaccinated population. The *overall effects* are defined by the contrast in the average outcomes in the entire population where some individuals were vaccinated compared to the average outcomes of the entire population that did not receive vaccine.

**Figure 1:**
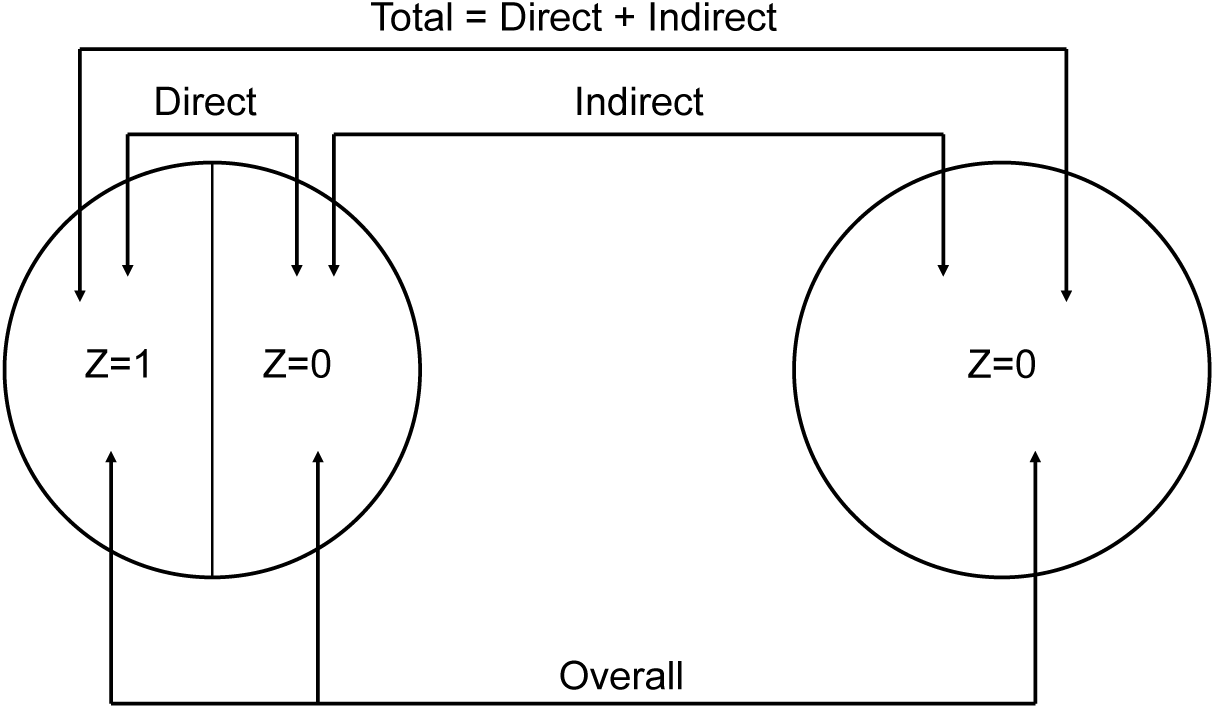
Study designs for dependent happenings. Two clusters, or populations, are considered under two different scenarios. In the scenario on the left, a certain portion of individuals in the cluster receive treatment, *Z =* 1, and the other portion of individuals receive control or nothing, *Z =* 0. In the scenario on the right, everyone receives control or nothing. The direct, indirect, total, and overall effects of intervention are defined by the indicated contrasts (adapted from Halloran and Struchiner 1991, 1995). The effects have recently been called something else in the economics literature: direct effect = value of treatment; indirect effect = spillover effect on the non-treated; overall effect = total causal effect; total effect = intention to treat effect (Baird et al 2017).

In the simple Figure 1, we have not distinguished trial participants from nonparticipants. On the population in which some individuals were vaccinated in Figure 1, some of the unvaccinated may be in the control arm of the trial, and some may not be in the trial. The indirect effects may reduce the incidence of infection for others within or outside the trial, invalidating the simple assumption that the incidence before the trial will be similar to the incidence in those trial participants who do not receive the intervention. For complex behavioral interventions to reduce infectious disease transmission, such as education programs designed to encourage sexual abstinence to reduce transmission of infections, those who receive the intervention and change their behavior may cause others in the population to change their behaviors in ways that will also affect the risk of participants in the trial. Due to such effects, the mean incidence in each group may depend in complicated ways on the intervention and trial design, which can be addressed with simulations.

### 1.2 Measuring effects beyond the individual level

While the indirect effects discussed above can complicate the design and analysis of intervention trials, measuring them may be of scientific interest beyond, or even instead of, measuring the direct effects of the intervention in protecting individual recipients. Establishing that vaccination provides population-level effects that go beyond the direct effects in the vaccinated can have important consequences for public health policy. Some interventions, such as treatment-as-prevention of HIV, and transmission-blocking malaria vaccines, have only indirect effects. In Figure 1, the overall effect of an intervention is often the quantity of greatest interest for policy makers, as it summarizes the public health consequences of the choice of intervention strategy if adopted in a population [2]. Establishing that vaccination produces indirect effects in the unvaccinated can make a vaccination strategy more cost-effective. The expected size of these population-level effects depends not only on the size of the direct effect, but also factors related to the transmission of the disease in the population and the distribution of the intervention. Thus, additional tools that account for these interactions may be required to design studies to measure them.

If evaluating population-level effects of interventions, such as the total, indirect or overall effects, is of interest, then a cluster-randomized study will generally be the design of choice [2]. In simple cluster-randomized studies, the clusters are randomized to intervention or control. In two-stage randomized studies, clusters are randomized to one of several possible levels of coverage, also called saturation, of the intervention (possibly zero, or pure control), then individuals within the clusters are randomized to receive or not receive the intervention with probability equal to the coverage assigned to the cluster [4, 5].

Simulations can examine the properties of different types of cluster-randomized designs, whether parallel, (all randomized at beginning to intervention or control), stepped wedge (the order in which clusters receive intervention is randomized before the trial) [6], or something else. For example, the ring vaccination trial of an Ebola vaccine compared outcomes in rings of contacts and contacts of contacts around a detected case and randomized each ring to receive either immediate or delayed vaccination [7, 8]. For each type of design, simulations can answer questions such as: what is the required sample size with this design, given the likely degree of transmission during the trial [9]? What is the optimal choice of cluster size versus number of clusters? What are optimal coverage (saturation) levels across clusters [4]? Simulations can also compare different types of designs, clarifying the tradeoffs among these designs in power and bias for estimating various quantities of interest. Recently, simulations were used to design a stepped wedge cluster-randomized study of the effectiveness of adding solar-powered mosquito trapping systems to standard malaria interventions, examining different methods of temporally introducing the intervention across an island [10].

Simulations can also help predict potentially harmful indirect effects. In an intervention where women are encouraged to refuse sexual acts with men, other women in the population, either study participants or nonparticipants, might be sought out by the men refused by women in the trial, known as spillover effects [4] or displacement [11]. If displacement occurs, the incidence of HIV may be higher in the women not benefitting from the intervention than it would have been if the trial had not occurred. In malaria interventions, when some individuals use bednets, the individuals not using bednets may be bitten more often and have increased incidence of malaria. Alternatively, with insecticide-treated bednets, mosquitoes killed because one participant in the intervention arm of a trial uses a treated net, may then fail to bite a person who is in the control arm of the trial, reducing the risk in the control group, thus reducing power. These spillover effects are part of the dynamic process that can be included in simulations, and thus accounted for in the design phase.

### 1.3 Transmission as a cause of overdispersion

Not only the average risk, but also the variability in risk among individuals receiving a particular intervention may be hard to predict in the infectious disease context. Classical trial sample size calculations rely on simple assumptions about variability for independent events which may be invalid because they do not account for complex social and sexual networks. Because of the dynamics of the transmission process and random events, a group, such as a village or hospital, within an arm of a trial may have many more or many fewer cases than the average value. The amount of this variability may depend on how the trial is designed, for example whether individual persons or groups of persons are randomized to trial arms, and how the intervention is rolled out over time in the trial. For example, in HIV prevention trials, one needs to account for stochastic variation in the spread of HIV from overlapping sexual networks and heterogeneity in biological and behavioral risk factors in addition to the usual variability in standard study design [12]. Simulations can take into account different sources of variation as well as do sensitivity to unknown sources of variation in calculating sample sizes.

### 1.4 Combinations of interventions

Combinations of interventions in infectious diseases may have interactions at the population level that are difficult to predict or to express in simple equations. When different single interventions or combinations of interventions are being considered, simulations can be used to explore possibly synergistic effects on transmission and outcomes relevant for trial design. Boren et al [12] used simulations to estimate the effect size expected from each of four HIV-prevention interventions if they were implemented individually or in varying combinations in a South African population. These effect-size estimates could be the basis for sample-size calculations in a cluster-randomized trial and for evaluating which interventions to test first. Here, assumptions could be made about the effect of each intervention on its own on an individual’s risk of contracting HIV, but simulations were necessary to understand how these interventions affected transmission in the population, via indirect effects, as well as to understand how the effects of the different interventions combined at the population level.

### 1.5 Heterogeneities in hosts and pathogens

These complexities of trial design can be compounded by additional sources of heterogeneity. Individuals may differ dramatically in both their exposure to infection and their responsiveness to the intervention. Trial planners may have little advance knowledge about the distribution of these sources of variability. Interactions between different pathogen strains can create further variability in risk of the outcome that is difficult to incorporate into simple equations. An example is when a vaccine, such as pneumococcal vaccine, protects against some but not all strains of the pathogen, and individuals who acquire one strain are protected against acquiring another strain, potentially making a vaccine seem more or less efficacious than it is [13]. In addition, infectious diseases have epidemics that have an intrinsic tendency to grow as more people become infected, increasing the risk to others, and eventually to contract as previously susceptible people become infected and immune. Thus the inputs into sample size calculations are a moving target in infectious diseases, sometimes moving quite fast and varying spatially. Sometimes the actual quantities that can be measured from the data observed in a trial are nonlinear functions of the biological efficacy of a vaccine or drug [13, 15], which may be the quantity of most direct interest or may be the quantity considered most likely to be transportable to populations beyond that in which the trial was conducted.

Simulations can help capture the impact of heterogeneity in the trial population on power and on the size of the effect being measured. For example, heterogeneity in the exposure or susceptibility of trial participants to infection can bias vaccine efficacy estimates toward the null, but simulations showed that under certain assumptions these biases can be avoided by accounting for variation in frailty in the analysis [16]. Simulations taking strain-to-strain interactions into account have demonstrated that even if estimation of heterogeneous protection of each component against pneumococcal carriage fails, unbiased and interpretable estimation of summary measures of vaccine efficacy may still be possible from the observed data [17].

### 1.6 Improving trial logistics and ethics

Simulations can help identify key elements determining the potential for success of a specific trial, beyond the choice of sample size and outcome measures. These include speed of case ascertainment, test results, or other time-sensitive processes that affect the likelihood of success. Ethical or logistical reasons may motivate the choice of design, particularly for trials in the emerging infectious disease or outbreak setting. The debate over trials of Ebola vaccines during the 2014-5 epidemic in West Africa illustrates that, especially in emergency situations, these considerations may place competing pressures on study design [18]. For example, in the Ebola ring vaccination trial in Guinea, in the face of declining transmission, if cases were not ascertained quickly enough, and the contacts of the cases and the contacts of contacts not found in a timely fashion, then the trial likely would not have been feasible. The ring-vaccination cluster-randomized trial ultimately demonstrated the effectiveness of one Ebola vaccine [7, 8]. The lesson was learned in Ebola to go where the transmission is or where it is expected to be. If vaccination had been allocated randomly in the population, the trial would not have had sufficient power for a conclusive result. Simulations are currently being used to help identify potential sites and study designs for Zika vaccine trials.

Besides being good scientific practice, designing a trial with adequate power is also an ethical requirement. Underpowered studies create burdens on participants without providing a high degree of assurance that the scientific results will be valuable [19], while unduly large studies place burdens on too many participants without significant extra benefits from the larger sample size. Poorly considered design runs the risk of discarding interventions that might be useful. Simulations can help find the design that is optimal in terms of either power, speed, or number of deaths/cases averted, that also fulfills ethical requirements for respecting human dignity and other ethical tenets. For example, a cluster-randomized stepped wedge trial was proposed to vaccinate frontline health care workers with a candidate Ebola vaccine as quickly as logistically feasible, but to randomize the order in which each treatment unit received vaccination to allow randomized evaluation of vaccine efficacy. Simulations found that an individually randomized controlled trial that prioritized the highest risk treatment units first indeed would have greater statistical power and speed to a definitive result, avoiding more total health care infections, than the proposed stepped wedge design [20]. This kind of work has a long history in cancer treatment trials [21], but has been less used in infectious diseases [22, 23]. The expected value of the information from a trial for decision-makers relative to its cost can also be easily calculated using simulation models with an economic component [24]. In the future, interactions among trial designers, epidemiologists, infectious disease modelers, and ethicists may help to specify ethical desiderata for trials and the designs best suited to accomplish these while maintaining ethical treatment individuals and study validity.

### 1.7 Interpreting Results of a Trial

After a trial, simulations can be helpful in interpreting puzzling or unexpected trial results. A cluster-randomized trial was conducted to compare the effectiveness of isoniazid preventive therapy given on a community-wide basis to standard of care on TB in gold miners in South Africa [25]. Although pre-trial mathematical modeling suggested that the intervention had unusually high potential for TB control [26], the intervention trial demonstrated no effectiveness on TB. This was thought to be due to lower than expected uptake. Post-trial modeling took advantage of data from the trial and demonstrated that even with optimal uptake, a combination of interventions would be required to greatly reduce TB incidence [27].

Trials assessing the effect of sexually transmitted disease treatment on HIV incidence in Rakai, Uganda, and Mwanza, Tanzania, had differing results, with a larger effect for the syndromic treatment intervention in Mwanza than for the mass treatment intervention in Rakai [28, 29]. Simulation research done in the early 2000s suggested that population differences in sexual behavior, curable sexually transmitted disease rates, and HIV epidemic stage could explain most of the contrast [30, 31].

Not all phenomena can be explained with traditional epidemic models. Simulation studies can help separate plausible from implausible hypotheses, evaluation of which can be incorporated into the design of later trials and field studies. Multiple randomized control trials showed oral cholera vaccine to be safe and effective. Detailed spatial analysis of the results of cholera vaccine trials in Matlab, Bangladesh, and Kolkata, India, revealed something peculiar. The strength of indirect effects increases with coverage faster in unvaccinated than vaccinated individuals [32, 33, 34], a result not yet explained by modeling. In this and similar instances, perhaps further biological understanding will be needed.

## 2 Simulation Approaches for Infectious Disease Trial Design

An increasingly popular approach to dealing with these challenges in infectious disease intervention trial design is to employ computer simulations of the trial in the setting of ongoing disease transmission. As illustrated in Figure 2, such simulations translate assumptions about the effect of an intervention on an individual’s risk, via a mechanistic dynamic model of the disease transmission process and the intervention in the trial setting, into predictions about the magnitude and variability of disease incidence in each arm of the trial. These simulations model explicitly the process of transmission, including such factors as population and contact network structure, natural history of disease and infectiousness, the phase of the epidemic, assumptions about the direct effects of intervention, and, for complex interventions, the timing and logistics of intervention in the trial.

**Figure 2:**
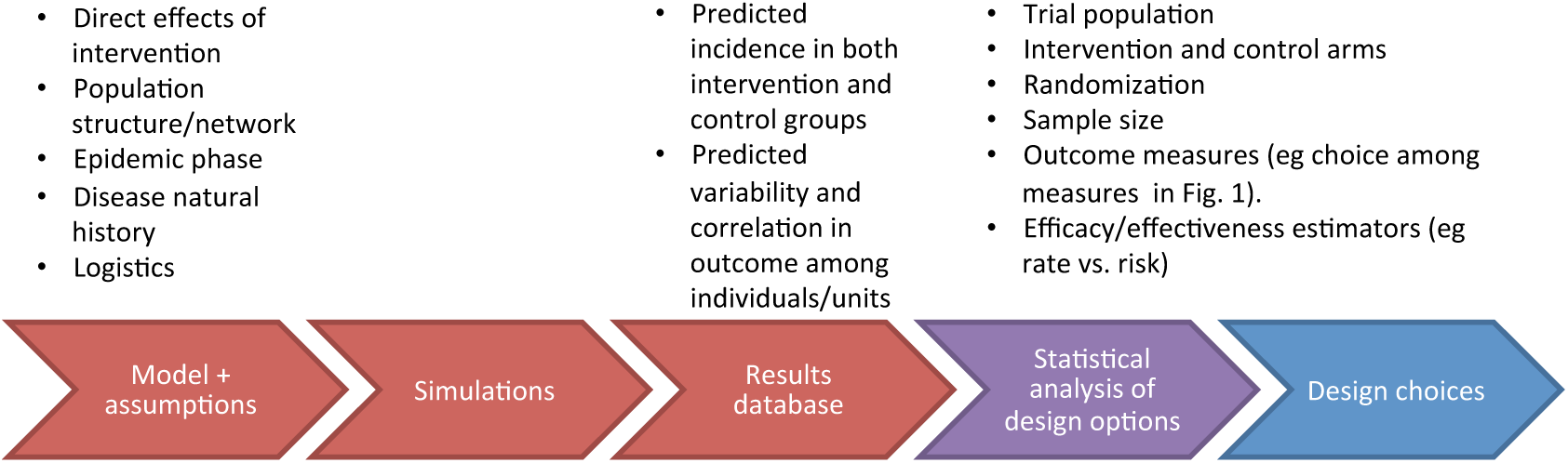
Illustration of simulations used for design and analysis of infectious disease intervention trials.

### 2.1 Simulating a trial

Within this synthetic population, a subset of individuals is selected as a trial population, and simulations of the planned trial are then conducted to generate observations similar to those that could be made in the trial, incorporating the stochasticity, or randomness, inherent in the intervention, the transmission process during the trial, and the observation process. The randomness is generated by figuratively flipping a biased coin in the computer for every possible event. Each single stochastic simulation produces data similar to that which might be observed in one trial. The simulation results can be stored in a database. From each simulated trial, the effect measures of interest can be estimated using a proposed method of statistical data analysis. The mean estimate over the set of simulations under given assumptions allows assessment of whether the trial will provide an unbiased estimate of the quantity of interest, while variability in the estimates produced over a series of simulated trials provides a measure of the likely variability. Sample size and power calculations can be derived directly from the variability of output of many simulations. Alternatively, intensity and patterns of variability of disease incidence within the simulated trial can be used as inputs to conventional formulas to calculate the effect size that will be detected in the trial and the sample size required to detect such an effect size given the level of variability.

By changing the assumptions of the simulation, it is possible to explore the sensitivity of the measured effect size or its variability to particular aspects of trial implementation, and thereby to design more efficient trials. By analyzing simulation results with different statistical methods, it is possible to identify potential biases in the analysis that arise from the transmission process.

### 2.2 Choice of simulation approach

A key question is what level of detail is required in the simulation. One can go from full calibration to a country, such as South Africa [35], to simple simulations with dynamics but little explicit demographic structure [12]. Simplicity needs to be balanced with sufficient detail such that the bias in the results is minimized. One needs to carefully weigh each additional complexity and consider whether the choice of intervention and study design is likely to be affected by it. If comparing analytical approaches only, then purely statistical models may be sufficient, as in [6]. In many contexts, however, dynamic mechanistic models are more useful. In some cases it may be important to consider even within-host effects.

Simulations for designing trials need to choose a contact network structure and critically examine that choice. The choice of network structure used to design an intervention trial can affect the predicted power and needed sample size for the trial. Simulations using networks can examine the effect of within-cluster structure on statistical power and sample size [36]. Simulations can similarly be used to examine the effect of between cluster mixing, that is, contamination across clusters, on the power of a study and bias of estimates. Incorporating information on network features can improve the efficiency of treatment effect estimation in cluster-randomized trials [37]. In general, simulation of trials on contact networks can help us understand under what circumstances the structure of these networks matters in trial design and when they can be ignored. At a finer level of detail, simulation can help identify network features that are relevant, what is a minimal set of such features that still yields gains in power, and how precisely such features would have to be measured in practice.

The choice of model needs to be matched to the question of interest. Stochastic, individual-based models and deterministic differential equation models can do different things related to trial design. Stochastic, individual-based models have the advantage of generating data that have statistical variation, and thus can better be used for trying out different statistical methods of analysis. They allow more detail in individual attributes and may also more accurately reproduce the population-level effects to be estimated. Differential equation models generally run much more quickly and can aid in studying the effects of different combinations of interventions on the outcome, and allow study of point estimates of effect measures. However, in general, they do not produce synthetic data that can be analyzed using the proposed statistical methods without an additional stochastic layer.

Concern about robustness of results to model misspecification is often raised. But advice that emanates from a model may still be qualitatively correct even if the model does not accurately characterize all aspects of the transmission system. For example, the advice in trade-offs for optimal design in estimating different effects of interventions in two-staged randomized studies [4] is model-assisted and is used to draw analytical insights that can be valid even if the model is imperfect.

One challenge to using simulations to design and interpret studies is the difficulty in having the appropriate data to inform all aspects of the simulations. This includes empirical epidemiological data about the natural history and transmission of the disease or treatment effect. Simulation can assess the impact of uncertainty or lack of knowledge on potentially important epidemic features. An advantage of using simulations after a trial is that a great deal of additional data will be available to inform the modeling.

In addition to increasing confidence in the choice of population and sample size, the process of designing, implementing and performing the simulations may provide other benefits, such as insight into potential biases in effect estimates, ways to account for heterogeneities in the trial population, calculations of quantities related to the ethics of trial design, and hints about choices in trial design beyond the sample size that may improve the probability of the trial’s success. Despite potential caveats, the relative cost of computations versus actually running a trial will be very low, while the potential gains in avoiding inconclusive studies are large.12

## 3 Conclusions

Expertise in infectious disease dynamics, statistical science, and simulation methods are required to adequately design and interpret trials for many interventions against current and emerging infectious diseases. Trial design should draw in an expert on infectious disease transmission dynamics, working alongside the statistician who is virtually always employed for study design. Indeed, an increasing number of investigators have both sets of skills. While simulation experiments cannot replace trials, nearly all trials would benefit from simulations, or at least from an exercise to write down the steps in such a simulation, which may in itself point out unexpected aspects of trial design. It is helpful in planning a trial to explicitly write down a timeline of the events in the trial for each case and consider possible variation in the sequence of these events, such as the diagram included in the design of the Ebola ring vaccination trial [7]. Clear communication of complex simulations, the assumptions underlying them and their limitations, is important. In addition to simulating trials prior to implementation, there is value in validating modeling approaches after a trial is completed to develop lessons learned for simulating the next trial. Over the long term, it could be useful to use a combination of simulations and practical experience to develop rules of thumb for adjustment that can be used as guides in smaller studies.

## Acknowledgments

This paper is the result of discussions at a workshop funded by the National Institute of General Medical Sciences MIDAS Center of Excellence grants U54 GM111274 and U54 GM088558.

## References

[1] R Ross. An application of the theory of probabilities to the study of *a priori* pathometry, Part 1. Proc R Soc Series A, 92:204–230, 1916.

[2] ME Halloran and CJ Struchiner. Study designs for dependent happenings. Epidemiology, 2:331–338, 1991.

[3] ME Halloran and CJ Struchiner. Causal inference for infectious diseases. Epidemiology, 6:142–151, 1995.

[4] S Baird, JA Bohren, C McIntosh, and B Özler. Optimal design of experiments in the presence of interference. Review of Economics and Statistics, forthcoming, 2017.

[5] MG Hudgens and ME Halloran. Towards causal inference with interference. J Am Stat Assoc, 103:832–842, 2008.

[6] MA Hussey and JP Hughes. Design and analysis of stepped wedge cluster randomized trials. Contemp Clin Trials, 28:182–191, 2007.

[7] Ebola #x00E7;a suffit ring vaccination trial consortium. The ring vaccination trial: a novel cluster randomised controlled trial design to evaluate vaccine efficacy and effectiveness during outbreaks, with special reference to Ebola. BMJ, 351:h3740, 2015.

[8] AM Henao-Restrepo, IM Longini, M Egger, NE Dean, et al. Efficacy and effectiveness of an rVSV vectored vaccine expressing Ebola surface glycoprotein: interim results from the Guinea ring vaccination cluster-randomised trial. Lancet, 386:857–66, 2015.

[9] MDT Hitchings, RF Grais, and M Lipsitch. Using simulation to aid trial design: Ring-vaccination trials. PLoS Negl Trop Dis, 11(3):e0005470, 2017. https://doi.org/10.1371/journal.pntd.0005470.

[10] M Silkey, T Homan, N Maire, A Hiscox, R Mukabana, W Takken, and TA Smith. Design of trials for interrupting the transmission of endemic pathogens. Trials, 17:278, 2016. doi: 10.1186/s13063-016-1378-1.

[11] AW McCormick, NN Abuelezam, T Fussell, GR Seage, and M Lipsitch. Displacement of sexual partnerships in trials of sexual behavior interventions: A model-based assessment of consequences. Epidemics, 17:doi: 10.1016/j.epidem.2017.03.007, 2017.

[12] D Boren, PS Sullivan, C Beyrer, SD Baral, L-G Bekkerd, and R Brookmeyer. Stochastic variation in network epidemic models: implications for the design of community level HIV prevention trials. Statistics in Medicine, 33:3894–3904, 2014.

[13] K Auranen, H Rinta-Kokko, and ME Halloran. Estimating strain-specific and overall efficacy of polyvalent vaccines against pathogens with recurrent dynamics from a cross-sectional study. Biometrics, 69:235–44, 2013.

[14] CJ Struchiner, ME Halloran, RC Brunet, JMC Ribeiro, and E Massad. Malaria vaccines: Lessons from field trials. Cadernos do Saúde Pública, 10(supplement 2):310–326, 1994.

[15] JJ O’Hagan, M Lipsitch, and MA Hernán. Estimating the per-exposure effect of infectious disease interventions. Epidemiology (Cambridge, Mass), 25(1):134–138, jan 2014.

[16] IM Longini and ME Halloran. A frailty mixture model for estimating vaccine efficacy. Appl Stat, 45:165–173, 1996.

[17] J Mehtää, R Dagan, and K Auranen. Estimation and interpretation of heterogeneous vaccine efficacy against recurrent infections. Biometrics, 72(3):976–985, 2016.

[18] National Academies of Sciences, Engineering, and Medicine. Integrating Clinical Research into Epidemic Response: The Ebola Experience. The National Academies Press, Washington, DC, 2017. https://doi.org/10.17226/24739.

[19] SW Lagakos and AR Gable. Challenges to HIV prevention - seeking effective measures in the absence of a vaccine. New England Journal of Medicine, 358:1543–1545, 2008.

[20] SE Bellan, JRC Pulliam, CAB Pearson, D Champredon, et al. Statistical power and validity of Ebola vaccine trials in Sierra Leone: a simulation study of trial design and analysis. Lancet Infectious Diseases, 15:703–710, 2015.

[21] DA Berry. Bayesian clinical trials. Nature reviews. Drug discovery, 5(1):27–36, January 2006.

[22] BS Cooper, MF Boni, W Pan-ngum, NPJ Day, PW Horby, P Olliaro, et al. Evaluating clinical trial designs for investigational treatments of Ebola virus disease. PLoS Medicine, 12(4):e1001815, 2015. doi:10.1371/journal.pmed.1001815.

[23] G Harling, R Wang, J-P Onnela, and V De Gruttola. Leveraging contact network structure in the design of cluster randomized trials. Clinical Trials (London, England), October 2016.

[24] JV Robotham, N Graves, BD Cookson, AG Barnett, JA Wilson, JD Edgeworth, R Batra, BH Cuthbertson, and BS Cooper. Screening, isolation, and decolonisation strategies in the control of meticillin resistant Staphylococcus aureus in intensive care units: cost effectiveness evaluation. BMJ, 343:d5694, 2011.

[25] GJ Churchyard, KL Fielding, JJ Lewis, L Coetzee, EL Corbett, et al. A trial of mass isoniazid preventive therapy for tuberculosis control. N Eng J Med, 370:301–310, 2014.

[26] KL Fielding, AS Grant, RJ Hayes, RE Chaisson, EL Corbett, and GJ Churchyard. Thibela TB: design and methods of a cluster randomised trial of the effect of community-wide isoniazid preventive therapy on tuberculosis amongst gold miners in South Africa. Contemp Clin Trials, 32:382–392, 2011.

[27] E Vynnycky, T Sumner, KL Fielding, JJ Lewis, AP Cox, et al. Tuberculosis control in South African gold mines: Mathematical modeling of a trial of community-wide isoniazid preventive therapy. Am J Epidemiol, 181(8):619632, 2015. doi: 10.1093/aje/kwu320.

[28] H Grosskurth, F Mosha, J Todd, et al. Impact of improved treatment of sexually transmitted disease on HIV infection in rural Tanzania: Randomised controlled trial. Lancet, 346:530–536, 1995.

[29] MJ Wawer, NK Sewankambo, D Serwadda, et al. Control of sexually transmitted disease for AIDS prevention in Uganda: a randomized community trial. Lancet, 353:525–535, 1999.

[30] RG White, KK Orroth, EL Korenromp, R Bakker, M Wambura, NK Sewankambo, et al. Can population differences explain the contrasting results of the Mwanza, Rakai, and Masaka HIV/sexually transmitted disease intervention trials?: A modeling study. J Acquir Immune Defic Syndr, 37:1500–13, 2004.

[31] KK Orroth, RG White, EL Korenromp, R Bakker, J Changalucha, JD Habbema, and RJ Hayes. Empirical observations underestimate the proportion of human immunodeficiency virus infections attributable to sexually transmitted diseases in the Mwanza and Rakai sexually transmitted disease treatment trials: Simulation results. Sex Transm Dis, 33(9):536–44, 2006.

[32] M Ali, M Emch, M von Seidlein, M Yunus, DA Sack, M Rao, J Holmgren, and JD Clemens. Herd immunity conferred by killed oral cholera vaccines in Bangladesh: A reanalysis. Lancet, 366:44–49, 2005.

[33] M Ali, D Sur, YA You, S Kanungo, B Sah, B Manna, M Puri, et al. Herd protection by a bivalent killed whole-cell oral cholera vaccine in the slums of Kolkata, India. Clin Inf Dis, 56:1123–1131, 2013.

[34] C Perez-Heydrich, MG Hudgens, ME Halloran, JD Clemens, M Ali, and ME Emch. Assessing effects of cholera vaccination in the presence of interference. Biometrics, 70(3):731–741, 2014.

[35] AW McCormick, NN Abuelezam, ER Rhode, T Hou, RP Walensky, et al. Development, calibration and performance of an HIV transmission model incorporating natural history and behavioral patterns: Application in South Africa. PLoS ONE, 9(5):e98272, 2014. doi:10.1371/journal.pone.0098272.

[36] P Staples, EL Ogburn, and J-P Onnela. Incorporating contact network structure in cluster randomized trials. Scientific Reports, 5:doi:10.1038/srep17581, 2015.

[37] P Staples, M Prague, V DeGruttola, and J-P Onnela. Leveraging contact network information in cluster randomized trials of infectious processes. arXiv.org, arXiv:1610.00039 [stat.AP], 2016.

